# *Caenorhabditis elegans* dauers vary recovery in response to bacteria from natural habitat

**DOI:** 10.1101/2020.04.01.020693

**Authors:** Louis T. Bubrig, John M. Sutton, Janna L. Fierst

## Abstract

Many species use dormant stages for habitat selection by tying recovery from the stage to informative external cues. Other species have an undiscerning strategy in which they recover randomly despite having advanced sensory systems. We investigated whether elements of a species’ habitat structure and life history can bar it from developing a discerning recovery strategy. The nematode *Caenorhabditis elegans* has a dormant stage called the dauer larva that disperses between habitat patches. On one hand, *C. elegans* colonization success is profoundly influenced by the bacteria found in its habitat patches, so we might expect this to select for a discerning strategy. On the other hand, *C. elegans*’ habitat structure and life history suggest that there is no fitness benefit to varying recovery, which might select for an undiscerning strategy. We exposed dauers of three genotypes to a range of bacteria acquired from the worms’ natural habitat. We found that *C. elegans* dauers recover in all conditions but increase recovery on certain bacteria depending on the worm’s genotype, suggesting a combination of undiscerning and discerning strategies. Additionally, the worms’ responses did not match the bacteria’s objective quality, suggesting that their decision is based on other characteristics.

## Introduction

Many organisms use developmentally-arrested dormant stages to endure harsh environments and/or disperse to better ones (***Baskin and Baskin, 1998***). Dormant stages must recover to resume growth but this transition is often irreversible and exposes the individual to new dangers (***Raimondi, 1988***). Therefore, individuals that assess local conditions and tie this information to their recovery can increase their fitness (***Keough and Downes, 1982***). Unsurprisingly, this has led to the evolution of a diversity of discerning strategies (***Baskin and Baskin, 1998; Johnson et al., 1997***). The cues that induce dormant stage recovery are tailored to the organism’s abiotic and biotic needs; the strategies can be as simple as measuring temperature (***Finch-Savage and Leubner-Metzger, 2006***) or detecting conspecifics (***Burke, 1986***) and as complicated as parsing out signals from whole communities. Coral larvae, for example, can differentiate between algal species growing in a prospective settlement site (***Harrington et al., 2004***). While many species develop these discerning strategies, other species seem to adopt an undiscerning strategy, recovering under all conditions, even poor ones (***Keough and Downes, 1982***). If these species have variable habitat qualities that impact their fitness, why aren’t discerning strategies being selected for?

One possible explanation is that discerning strategies only arise if they help organisms avoid bad habitats and find good ones. A dormant organism may ignore salient information about its environment if it has no capacity to act on it (***Raimondi, 1988***). Behavioral constraints, life history traits, and habitat structure may prevent the development of discerning strategies, even when they would seem useful at first glance. In this project, we investigated how the nematode *Caenorhabditis elegans* recovers from its dormant stage–the dauer (Fig. 1)–given that the species seems pulled in two opposite directions. On one hand, the dauer appears perfectly suited for a complex habitat recognition system. This dormant stage is carried by small invertebrates to new habitat patches that vary substantially in their quality with some patches being totally inhospitable due to their bacterial community composition (***Samuel et al., 2016; Kiontke and Sudhaus, 2006***). Bacteria can be good sources of food or deadly pathogens depending on the species (***Felix and Braendle, 2010; Samuel et al., 2016***) and *C. elegans* can certainly differentiate between them (***Johnson et al., 1997***), at least from a mechanistic standpoint. Recovering is an irreversible decision that affects fitness: dauers are hardy and long-lived but cannot reproduce (***Cassada and Russell, 1975; Klass and Hirsh, 1976; Ellenby, 1968***) while recovered worms can establish colonies but are vulnerable.

**Figure 1.**
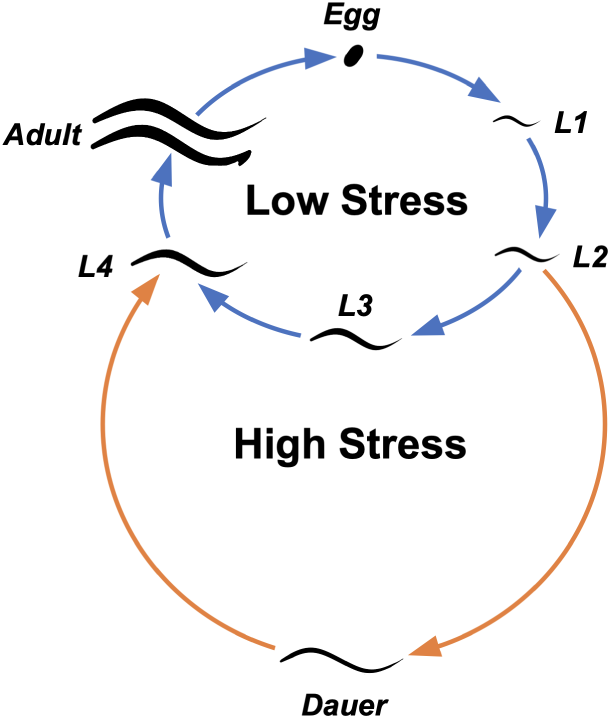
The life cycle of *C. elegans*. Newly hatched worms that sense high environmental stress become dauer larvae instead of the normal third larval stage (L3). Dauers that sense improving conditions can reenter the low stress cycle and continue to adulthood.

On the other hand, behavioral constraints and habitat structure may keep *C. elegans* from developing discerning recovery strategies. *C. elegans* dauers cannot control their invertebrate carriers and will be dropped off in bad habitats and good habitats alike. Unlike seeds which can stay put and ride out bad conditions for years (***Baskin and Baskin, 1998***), *C. elegans*’s natural habitats are ephemeral, rotting away in a matter of days (***Ferrari et al., 2017***). Unlike many marine invertebrates which can reject bad sites and move on to others (***Pawlik, 1992***), we have no evidence that *C. elegans* can do the same; the worms are likely stuck wherever they first arrive. External cues are only useful if they are actionable (***Raimondi, 1988***), so the worms’ lack of choice may lead them to ignore these cues in favor of simply recovering indiscriminately in the hopes of establishing a foothold.

We investigated how these opposing aspects of *C. elegans*’ ecology translate into recovery strategies by exposing dauers to a range of bacteria. We used four ecologically-relevant bacterial species isolated from *C. elegans*’ natural habitat (***Samuel et al., 2016***). We also sequenced the genomes of these four bacteria to facilitate future studies into natural worm-bacteria interactions. ***Samuel et al., 2016*** categorized each bacterial species based on *C. elegans* population growth and immune system activation. *Raoultella* sp. JUb54 and *Providencia* sp. JUb39 are considered “beneficial” because they support *C. elegans* population growth and do not activate the worm’s immune system. *Serratia* sp. JUb9 and *Pseudomonas* sp. BIGb0427 are “detrimental” because they are pathogenic and cannot support *C. elegans* populations. In addition to the natural bacteria, we included *Escherichia coli* OP50, the standard laboratory food which is not a natural food source (***Frezal and Felix, 2015***), and a control treatment with no food at all. To determine if *C. elegans* exhibits intraspecific variation in dormancy recovery, we tested three different worm strains that are geographically and genetically distinct. N2, isolated in Bristol, is the *C. elegans* reference strain which has been used since the mid 1900s. CB4856 is a very distant relative isolated in Hawaii. JU1395 is a much more recent isolate taken from France in 2008. We exposed dauers to bacteria for three hours, after which we collected and scored them based on their recovery status. Our data suggest that *C. elegans* dauer recovery has elements of both undiscerning and discerning strategies: *C. elegans* dauers recover regardless of condition but enhance their recovery when detecting certain bacteria. Additionally, *C. elegans* exhibits intraspecific variation in its recovery behavior.

## Results

Observations are summarized in Table 1. Of the 19,071 worms observed in this project, 8384 (or about 44%) recovered from the dauer stage after a three hour exposure. Recovery was not evenly distributed among the worm strains. N2 worms recovered the least–about 34.4%–which is consistent with previous work on recovery in this strain (***Cassada and Russell, 1975***). CB4856 had a slightly higher recovery at 39.2% while JU1395 had a much higher recovery at 56.4% (Fig. 2). Additionally, there were some batch effects among the trials; the worms in certain trials had depressed or enhanced recovery across the board (Fig. A1).

**Table 1.** Summary of observations categorized by worm strain, bacterial treatment, and recovery status.

|  |  | Control | *E. coli* | *Raoultella* | *Providencia* | *Pseudomonas* | *Serratia* |
| --- | --- | --- | --- | --- | --- | --- | --- |
| **N2** | Total Worms | 654 | 808 | 980 | 921 | 987 | 1372 |
|  | % Recovered | 29.2% | 38.0% | 36.0% | 36.3% | 33.4% | 32.9% |
| **CB4856** | Total Worms | 1011 | 954 | 1258 | 895 | 896 | 1438 |
|  | % Recovered | 32.6% | 42.6% | 40.2% | 36.2% | 37.6% | 43.4% |
| **JU1395** | Total Worms | 1048 | 1031 | 1374 | 1125 | 1112 | 1207 |
|  | % Recovered | 50.6% | 52.5% | 56.8% | 66.7% | 53.3% | 57.7% |

**Figure 2.**
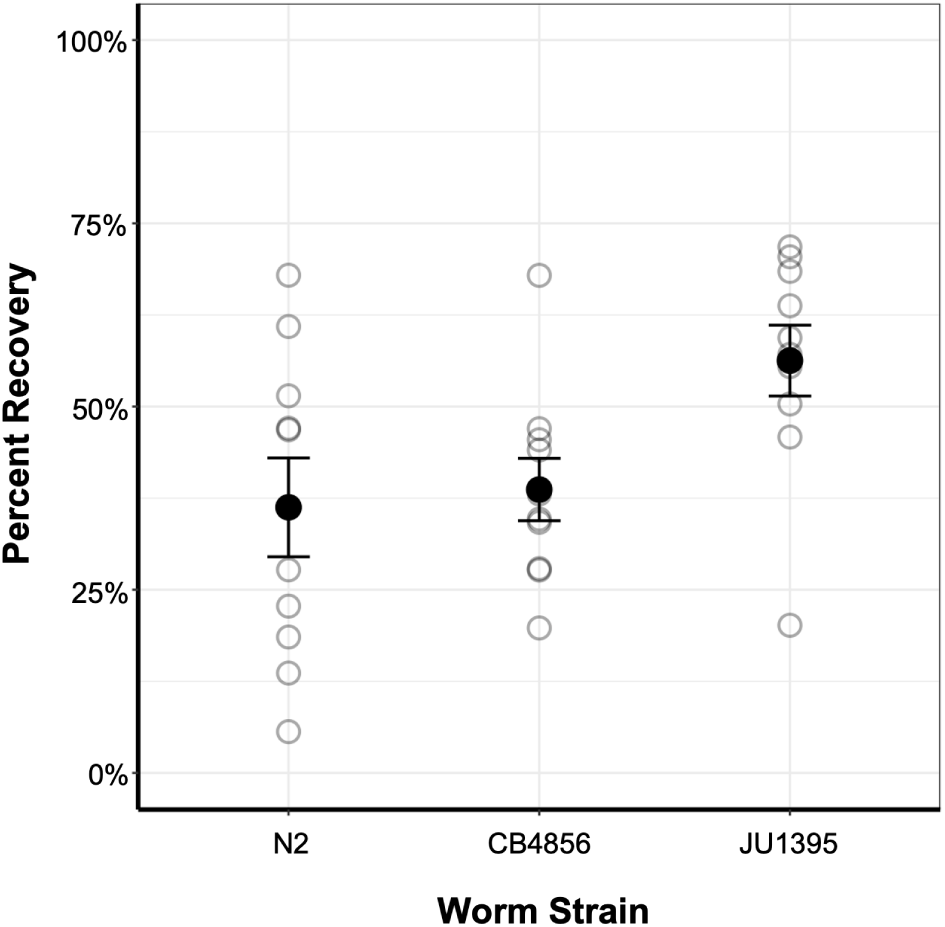
Mean recovery for the three worm strains. Faded points are average recovery values for each trial with all treatments combined. Error bars show standard error of the mean.

Worm recovery depended on bacterial treatment but also on which strain was detecting the bacteria, suggesting an interaction between these two variables (Fig. 3). N2 had broadly enhanced recovery on all beneficial bacteria with the highest mean recovery on *E. coli*. N2 also enhanced its recovery on the detrimental bacteria but only marginally. CB4856’s recovery was similar to N2’s but included an enhanced recovery on the detrimental bacterium *Serratia* sp. JUb9. JU1395 recovered the most on the beneficial bacterium *Providencia* sp. JUb39. JU1395’s recovery on *Serratia* sp. JUb9 was also very high, although this seems driven by one outlier during trial 2 in which JU1395’s recovery increased by a factor of 4.60.

**Figure 3.**
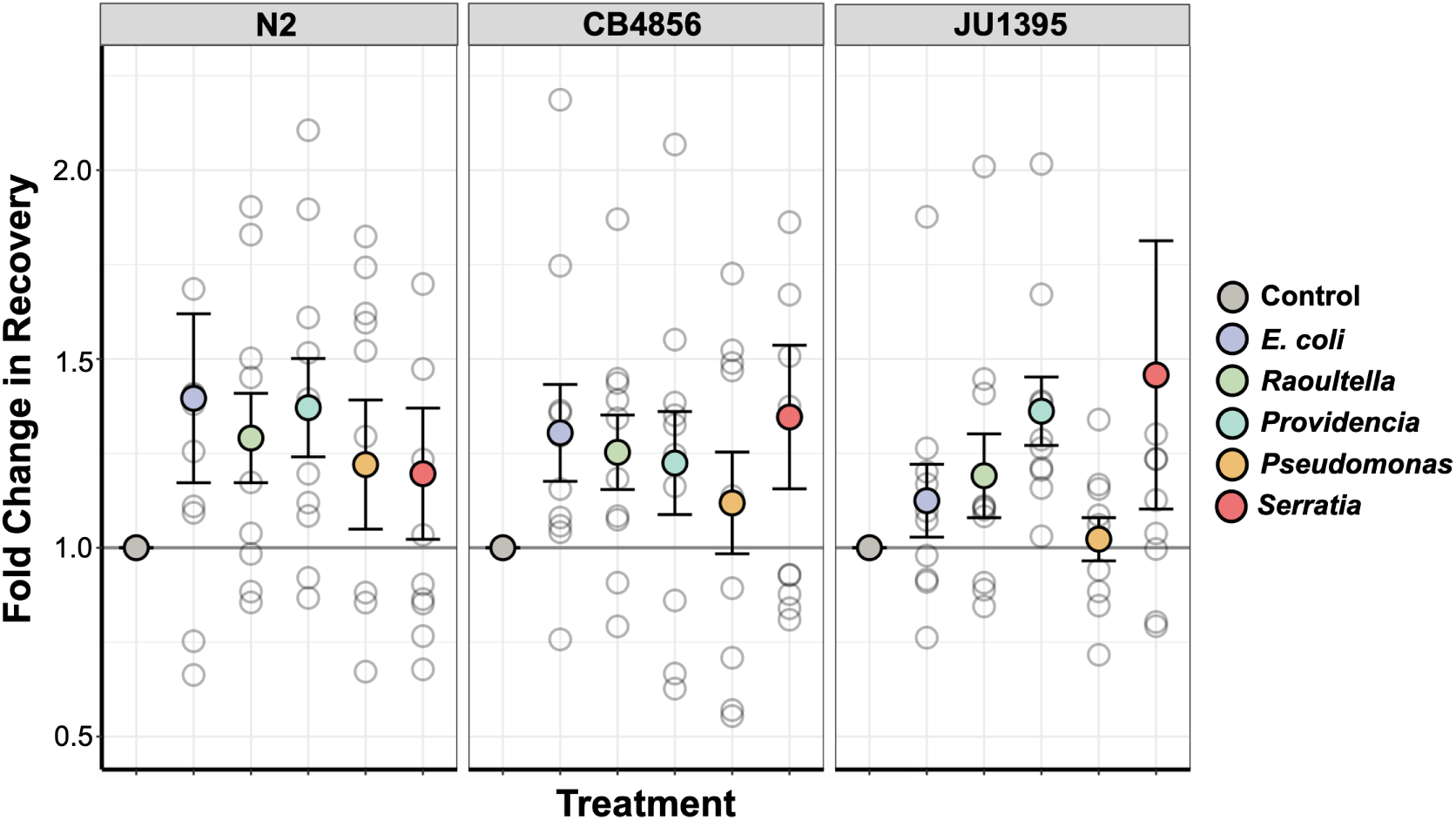
Fold change in recovery standardized by the percent recovered on the control of each trial. Cool colors represent beneficial bacteria and warm colors represent detrimental bacteria. Error bars show standard error of the mean. Five outlier points lie off the graph: N2 on *E. coli* OP50 has a value at 3.21; N2 on *Pseudomonas* sp. BIGb0427 has a value at 0.20; N2 on *Serratia* sp. JUb9 has a value at 2.46; CB4856 on *Serratia* sp. JUb9 has a value at 2.67; JU1395 on *Serratia* sp. JUb9 has a value at 4.60.

When categorizing the bacterial species, ***Samuel et al., 2016*** only performed worm growth assays using the N2 strain. We expanded this assay to include CB4856 and JU1395. We found that CB4856 and JU1395 grow no differently than N2 on the range of bacteria, so the categorizations established in ***Samuel et al., 2016*** hold. Worms on beneficial bacteria reached adulthood and produced eggs somewhere between 50 and 70.5 hours after they began feeding (Fig. A2). *Serratia* sp. JUb9 attracted and killed worms such that the population could not progress past the first few larval stages. *Pseudomonas* sp. BIGb0427 repelled worms, keeping them in the first larval stage (L1) or the dauer stage. A few individuals managed to reach adulthood on the *Pseudomonas* sp. BIGb0427 plates, but this was likely due to scavenging contaminants outside the lawn; the same phenomenon occurred on control plates that had no food.

### Statistical Analysis

Because recovering from dauer is a binary developmental choice, we built a logistic regression model to explore which variables affected a worm’s probability of recovering. The basic results of the model are shown in Table 2. The model uses the worm strain N2 and the control treatment as baselines. Odds ratios represent the fold-change in probability of recovering compared to the baseline. For example, any worm recovering on *E. coli* as opposed to the control has a 1.70-fold increased probability of recovering. Odds ratios for the remaining variables can be found in Table A1.

**Table 2.**
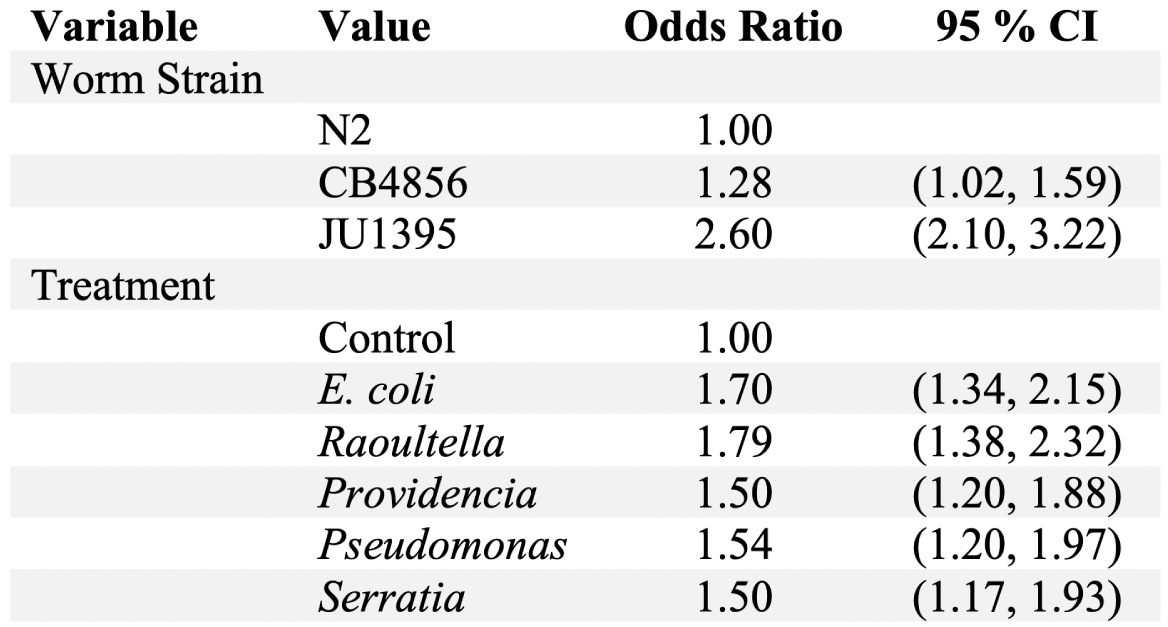
Estimated odds ratios for each value of the variables “Worm Strain” and “Treatment”.

Our model shows a significant interaction between “Worm Strain” and “Treatment”. This means that the odds ratios listed under “Treatment” in Table 2 should vary with worm strain. Table 3 shows the amounts by which they are adjusted, as well as the resulting odds ratios. Because N2 is the baseline worm strain and the control is the baseline treatment, N2 needs no adjustments, nor do any of the controls. The adjustments are made to the original odds ratios by simple multiplication. For example, a worm’s probability of recovery is predicted to increase 1.70-fold when exposed to *E. coli*. CB4856, however, is 0.92 times less likely to recover on *E. coli* than N2, the baseline worm strain. Therefore, CB4856’s recovery on *E. coli* is actually only 1.56-fold higher than its recovery on the control.

**Table 3.** Odds ratios of treatments adjusted due to interactions between “Worm Strain” and “Treatment”.

| Worm Strain | Treatment | Odds Ratio (without interaction) | Odds Ratio Adjustment | Odds Ratio (with interaction) |
| --- | --- | --- | --- | --- |
| N2 | Control | 1.00 |  | 1.00 |
|  | *E. coli* | 1.70 |  | 1.70 |
|  | *Raoultella* | 1.79 |  | 1.79 |
|  | *Providencia* | 1.50 |  | 1.50 |
|  | *Pseudomonas* | 1.54 |  | 1.54 |
|  | *Serratia* | 1.50 |  | 1.50 |
| CB4856 | Control | 1.00 |  | 1.00 |
|  | *E. coli* | 1.70 | 0.92 | 1.56 |
|  | *Raoultella* | 1.79 | 0.86 | 1.53 |
|  | *Providencia* | 1.50 | 0.79 | 1.18 |
|  | *Pseudomonas* | 1.54 | 0.90 | 1.38 |
|  | *Serratia* | 1.50 | 1.06 | 1.58 |
| JU1395 | Control | 1.00 |  | 1.00 |
|  | *E. coli* | 1.70 | 0.75 | 1.28 |
|  | *Raoultella* | 1.79 | 0.85 | 1.52 |
|  | *Providencia* | 1.50 | 1.32 | 1.98 |
|  | *Pseudomonas* | 1.54 | 0.74 | 1.14 |
|  | *Serratia* | 1.50 | 1.09 | 1.64 |

### Bacteria Sequencing

The results of our sequencing are shown in Table 4. We found that all of the wild bacteria except *Providencia* sp. JUb39 were closely related to previously reported genomes, albeit in unnamed species. We also found that the isolate JUb54, which was called *Enterobacter* sp. JUb54 in ***Samuel et al., 2016***, actually belonged to the genus *Raoultella* which is reflected in this article. Interestingly, *Serratia* sp. JUb9–which was found associated with *C. elegans* in France (***Samuel et al., 2016***)–is closely related to an isolate that was found in *C. elegans* habitats in Germany (Accession number: CP023268).

**Table 4.** Summary of information about sequenced bacteria.

| Species | Category | Genome size | Number of contigs |
| --- | --- | --- | --- |
| *Escherichia coli* OP50 | Beneficial | 4,616,404 | 1 |
| *Raoultella* sp. JUb54 | Beneficial | 5,422,632 | 1 |
| *Providencia* sp. JUb39 | Beneficial | 4,340,164 | 2 |
| *Pseudomonas* sp. BIGb0427 | Detrimental | 5,864,124 | 7 |
| *Serratia* sp. JUb9 | Detrimental | 5,108,081 | 1 |

### Dauer genes

*C. elegans* dauer entry and recovery are influenced by several well-characterized pathways including those underlying pheromone synthesis, guanylyl cyclase, TGF*β*-like, insulin-like and steroid hormone synthesis (***Girard et al., 2007***). Since the three worm strains responded differently to the range of bacteria, we sought to characterize molecular polymorphisms in these conserved dauer-controlling pathways. N2 and CB4856 already had sequenced and assembled genomes (***Kim et al., 2019***), so we sequenced JU1395’s genome to allow for comparisons between the three strains. The assembled sequence was 103,053,620 nucleotides in 161 contiguous pieces. We used the software BUSCO to estimate the completeness of the assembled sequence by searching for a set of 3,131 genes thought to be conserved across nematodes (***Seppey et al., 2019***). We identified 98% of these genes in our assembled sequence with 97.4% found in complete single copy, 0.6% duplicated, 0.5% fragmented and 1.5% missing. For reference, the N2 *C. elegans* assembled genome sequence has 98.5% of this 3,131 gene set with 98% in single copy, 0.5% duplicated, 0.3% fragmented and 1.2% missing.

We aligned 113 *C. elegans* transcripts from 67 dauer-associated genes to the assembled CB4856 and JU1395 sequences. Neither genome has been fully annotated for protein-coding genes and we used these alignments to measure polymorphisms and potential divergence in genes underlying these pathways. We identified relatively few polymorphisms in these sequences in JU1395 and CB4856. For example, there were only 18 polymorphisms in 9 genes between N2 and JU1395 and 46 polymorphisms in 15 genes between N2 and CB4856. The full list of dauer-associated pathways, genes and polymorphisms is given in the appendix. These polymorphisms are interesting targets for future studies investigating the genetic basis of the worm-microbe interactions.

## Discussion

When habitat quality affects an organism’s fitness, we expect natural selection to align an organism’s recovery with habitat quality. In the case of *C. elegans*, variation in habitat quality might select for worms that can differentiate between bacteria, a key determinant of establishment success. However, *C. elegans* disperses via a carrier and cannot choose its habitat; modulating dauer recovery might not provide worms with any advantage (***Raimondi, 1988***). In this case, the fittest strategy could be one of high rapid recovery across the board to outcompete other colonists. Our data is consistent with both of these hypotheses.

All three worm strains recovered substantially in all treatments–even in the absence of food–which suggests that some level of recovery is guaranteed, regardless of habitat quality. This supports the hypothesis in which *C. elegans* cannot choose its habitat and recovers no matter what. Presumably, worms that try to colonize a bad habitat have higher fitness than worms that refuse to try at all (***Johnson et al., 1997***). The basal level of recovery depended on the worm strain. N2 has the lowest basal recovery of the three strains. Interestingly, N2 is also reluctant to enter the dauer stage in the first place (***Lee et al., 2019***). CB4856 has a similar recovery as N2 despite their large genetic divergence. JU1395 has the highest recovery by far. These differences may result from variation in conserved dauer-controlling pathways. We found that the three strains have several polymorphisms in key dauer genes. For example, JU1395 has a polymorphism in *daf-22*, a gene involved in dauer pheromone synthesis (***Golden and Riddle, 1985***), while N2 and CB4856 have identical *daf-22* sequences. Determining these polymorphisms’ functional impact–if any–can be addressed in future work using the genetic tools available in *C. elegans*. From an evolutionary point of view, differences between the strains could reflect varying levels of acceptable risk. Some conditions, such as consistently high levels of pathogens, may favor more cautious strategies with slower recovery while other conditions select for a faster response. Strategies may also diverge when different strains regularly co-occur in the same habitat. A strain that frequently encounters a more cautious strain could benefit by recovering rapidly and establishing early. Timing developmental decisions to beat out other strains is not unheard of in nematodes; strains of the related nematode *Pristionchus pacificus* intentionally drive other strains of the same species into the dauer stage to stop them from feeding (***Bose et al., 2014***).

Dauer recovery differs among the bacterial treatments which is evidence for a more discerning strategy. Interestingly, the species does this in a way that is still consistent with the undiscerning strategy; no response is lower than the control but some bacteria can enhance recovery. Recovery will always occur, even in bad conditions, but can be accelerated upon detecting good conditions. What *C. elegans* interprets as “good,” however, is much more complicated than we had assumed. The worms’ responses do not simply reflect the objective quality of the bacteria. The most favorable bacteria–that is, the one which elicited the greatest response–differs with worm strain. N2 responds highly to *E. coli* OP50 and so does CB4856, but CB4856 also responds highly to the detrimental bacterium *Serratia* sp. JUb9. In contrast, JU1395 shows little response to *E. coli* OP50 but strongly responds to *Providencia* sp. JUb39. These results indicate a lack of matching between recovery and a bacterium’s objective quality. For instance, we demonstrated that *Serratia* sp. JUb9 rapidly kills all three worm strains and does not support growing populations. Despite this, CB4756 and JU1395 unexpectedly have enhanced dauer recovery on the bacterium even though the newly recovered population will fail to grow on it. Similarly, *Providencia* sp. JUb39 is objectively a nutritious food source but CB4856 has reduced recovery on it.

This lack of matching between food quality and response could have several explanations. Perhaps imperfect matching stems from the novelty of that food source. Certain combinations of worm strain and bacteria may never occur in nature or have occurred recently enough that selection has not had time to act (***Chew, 1977***). Imperfect matching could also occur when odorants are shared across many bacterial species, so selection on one worm-bacteria response spills over into other responses. It is also possible that worms can glean information about the bacterial community as a whole from interactions with individual species. Perhaps the presence of a specific bacterium in a community signals overall community health, substrate composition, or age of the patch (***Johnson et al., 1997***); some species of coral, for instance, deduce their depth by sensing the composition of nearby bacterial communities (***Webster et al., 2004***). Finally, bacteria may release odorants to specifically manipulate bacteriovore behavior. Bacteria may be under selection to evade detection or, in the case of pathogens, to attract vulnerable hosts. Dauer behavior is known to be manipulated by at least one non-nematode organism, the beetle *Exomala orientalis* (***Cinkornpumin et al., 2014***), so manipulation by bacteria is certainly feasible. Interestingly, *Serratia marcescens*, a congener of *Serratia* sp. JUb9, is strongly attractive to *C. elegans* despite its high pathogenicity (***Zhang et al., 2005; Pradel et al., 2007***), an observation that has puzzled many researchers.

Our results demonstrate that *C. elegans* dauers modulate their recovery based on the bacteria they detect in their new habitat. If these differences in recovery result from selection, this suggests that tying recovery to external cues still provides some kind of fitness benefit, even when the habitat structure bars dormant stages from dispersing to a better habitat in time or space. Perhaps the variety of strategies results from finer-scale fluctuations in habitat quality over the course of the rotting process. Additionally, conspecifics that frequently co-occur could maintain divergent strategies that vary in their levels of acceptable risk or other characteristics. In conclusion, behavioral strategies do not simply evolve in response to strong environmental pressures. A full understanding must take into account an organism’s ecological context, habitat structure, and life history, all of which contribute to the evolution of dormancy recovery strategies.

## Methods and Materials

### Worms and bacteria

The strains of *C. elegans* used for this project were N2, CB4856, and JU1395, which were received from the Caenorhabditis Genetics Center (CGC). N2 is the standard laboratory strain which was isolated in Bristol, UK in 1951 but not frozen until 1969. CB4856 was isolated in Hawaii in 1972 and JU1395 was isolated in Montsoreau, France in 2008.

*E. coli* OP50 was also received from the CGC. The four wild bacteria were all isolated from different sites in France between 2004 and 2009 (***Samuel et al., 2016***). *Providencia* sp. JUb39 and *Raoultella* sp. JUb54 were taken from rotting apples and *Serratia* sp. JUb9 was found in compost. These three species were acquired from Marie-Anne Félix at Institute of Biology of the Ecole Normale Supérieure (IBENS). *Pseudomonas* sp. BIGb0427 was isolated from the rotting stem of a butterbur plant and was acquired from Buck Samuel at Baylor College of Medicine. All worms and bacteria were frozen at −80 °C and aliquots thawed for each experimental replicate.

### Setting up experimental plates

Approximately three weeks before the experiment, worms of each strain were thawed and placed on 100 mm *E. coli*-seeded Nematode Growth Medium (NGM) plates (***Stiernagle, 2006***). These worms were incubated at 20 °C and expanded to seven plates per strain over the course of six days. The original thaw plates were discarded and the remaining six plates per strain were washed with water and the worms bleached using standard laboratory protocols to limit contamination (***Stiernagle, 2006***). Bleached eggs hatched overnight on a rocker at room temperature. The next day, hatched worms were placed onto six new *E. coli*-seeded NGM plates per strain. The worms were incubated at 20 °C for two weeks to induce dauer formation via starvation and overcrowding.

Experimental plates were 100 mm standard NGM plates. Three of these plates were used for the control treatment and contained an addition of 0.1% ampicillin, a broad-spectrum antibiotic used to prevent bacterial growth. Plates were assigned random number IDs to blind the experiment and ensure unbiased counting later on. Five bottles of 50 mL Luria Broth were inoculated with each of the five bacterial species and a sixth control bottle remained sterile. All bacteria were incubated overnight with *E. coli* at 37 °C and the other bacteria and the control at 25 °C.

The next day, bacterial absorbances were measured with a spectrophotometer and used with the equations in Table A2 to estimate the bacterial density in each broth. The eighteen experimental plates were seeded in six groups of three, one group per treatment. 5×10^7^ CFU of each bacterial species were deposited onto the plates and water added to bring the final volume up to 500 *μ*L to ensure even spreading. For the three control plates, the volume of sterile broth deposited was equal to the largest volume of bacteria added for that replicate. The liquid was then spread in an even lawn across the plate and let dry in a vent hood.

After two weeks of starvation, worms were washed off of their plates and treated with 1% sodium dodecyl sulfate (SDS) on a rocker table for 30 minutes. This treatment kills all worms except those in the dauer stage (***Cassada and Russell, 1975***). The worms were washed with water four times to remove the SDS and the final volume reduced to about 2 mL. Three aliquots of a 1:100 dilution of these worms were scanned for live worms to estimate live dauer density in the undiluted tubes. 2000 dauers were then deposited in the center of experimental plates which were air dried in a vent hood and then stored at room temperature. The total time of exposure from worm deposition to worm removal was three hours.

### Worm counting

The volume of worms placed in the center of experimental plates also contained the bodies of worms killed during the SDS wash, but most of the live worms explored the rest of the plate during the three-hour exposure. This central spot was cut out of the agar to leave only worms that were live at the time of deposition. Worms were then washed off each experimental plate, treated with 1% SDS for 30 minutes, and then washed four times with water to remove excess SDS. Ten 20 *μ*L aliquots per experimental plate were spotted onto an empty plate. Worms were then visually assayed for movement and given a maximum of three seconds to move before being declared dead. Moving worms were counted as having survived the SDS treatment, indicating that they had remained in dauer during the three hour exposure. Worms that did not move were counted as having been killed by the SDS wash, indicating that they had begun to recover from the dauer stage.

### Fecundity assay

Synchronized L1 larvae of all three worm strains were acquired by following standard bleaching protocols and hatching the eggs overnight (***Stiernagle, 2006***). Populations of L1 larvae were spotted onto 60-mm NGM plates with either no bacteria (the negative control) or 100 *μ*L of overnight bacterial cultures. These plates were maintained at room temperature and scanned periodically for the presence of eggs and the general health of the population. The assay was done in triplicate.

### Statistical Analysis

Logistic regression models were built in R version 3.6.2. Several models were compared using the likelihood-ratio test (***Hosmer and Lemeshow, 2000***). We retained all variables in the model because removing any of them significantly reduced the model’s fit. Because worm strains had unique patterns of recovery (Fig. 3), we also introduced an interaction term between the variables “Worm Strain” and “Treatment” and retained it in the model because it significantly increased the model’s fit.

### Bacterial genome sequencing

Overnight cultures of each bacterial isolate were grown at 25 °C, with the exception of *E. coli* OP50 which was grown at 37 °C; one mL of each culture was place in a 1.5mL tube and centrifuged to pellet the bacteria. Excess media was removed from the tube prior to gDNA extraction. Genomic DNA was extracted from each sample using a modified phenol-chloroform extraction (***Green and Sambrook, 2017***). One microgram of DNA from each sample was then prepared for multiplexed sequencing by attaching unique barcodes to each sample from the Oxford Nanopore Technologies (ONT) Native Bar-coding Kit (EXP-NBD104). Following ligation of the barcode sequences; the DNA from each sample was pooled in equimolar amounts and prepared for sequencing using the ONT Ligation Sequencing Kit (SQK-LSK109). The multiplexed sample was sequenced on a R9.4.1 flow cell using a GridION X5 platform. The sequence data were de-multiplexed and trimmed of barcode sequences using Porechop. Each genome was then assembled using Canu v1.8 (***Koren et al., 2017***).

### Nematode DNA Extraction, Sequencing and Analysis

*C. elegans* JU1395 worms were grown on several 100 mm NGM plates seeded with *E. coli* to achieve large population sizes. Worms were washed from the plates using M9 buffer, bleached using standard procedures, and the eggs hatched overnight (***Stiernagle, 2006***). We pelleted the worms, removed the supernatant, then flash-froze the pellet with liquid nitrogen. We then extracted the genomic DNA using a modified phenol-chloroform isolation (modified from ***Green and Sambrook, 2017***). gDNA fragments were size selected using the Short Read Eliminator Kit from Circulomics Inc. One microgram of DNA was used to create a sequencing library with the ONT Ligation Sequencing Kit (SQK-SK109) and sequenced on a R9.4.1 RevD flow cell using a GridION X5 platform. Adapter sequences were removed using Porechop and the genome assembled using Canu v 1.9 (***Koren et al., 2017***). The genome was polished using Illumina paired-end reads generated by the CeNDR project (***Cook et al., 2017***) and the Pilon software package (***Walker et al., 2014***). We used the BUSCO software v4.0.5 to estimate genic completeness with the nematoda_odb10 dataset (***Seppey et al., 2019***). We used the gmap-gsnap software (***Wu and Nacu, 2010***) to align the N2 dauer gene transcripts to the CB4856 and JU1395 genome sequences. Polymorphisms were identified with Samtools (***Li et al., 2009***) and Bcftools (***Li, 2011***).

## Data Accessibility

DNA sequence data generated during this project have been deposited with the National Center for Biotechnology Information under Bioproject PRJNA622250 for JU1395 and PRJNA622270 for the microbial samples.

Data is archived at XXXXXXXX.

## Competing Interests

The authors declare that there are no conflicts of interest.

## Author Contributions

LTB conceived the study, performed the experiment, analyzed the data, and wrote the manuscript. JMS performed DNA extraction, genome sequencing, genome assembly, and wrote these sections in the manuscript. JLF oversaw the experiment, performed genome assembly, gene comparisons, and wrote these sections in the manuscript.

## Acknowledgments

The authors would like to thank members of the Fierst lab, Amanda Gibson, Levi Morran, Jason Pienaar, and Jesualdo Fuentes-Gonzalez for helpful discussion and Chris Youssef for assistance in the lab.

LTB was supported by the Graduate Council Fellowship through the University of Alabama Graduate School.

Worms strains were provided by the CGC, which is funded by NIH Office of Research Infrastructure Programs (P40 OD010440). Wild bacteria were provided by Marie-Anne Félix and Buck Samuel.

### Appendix

**Figure A1.**
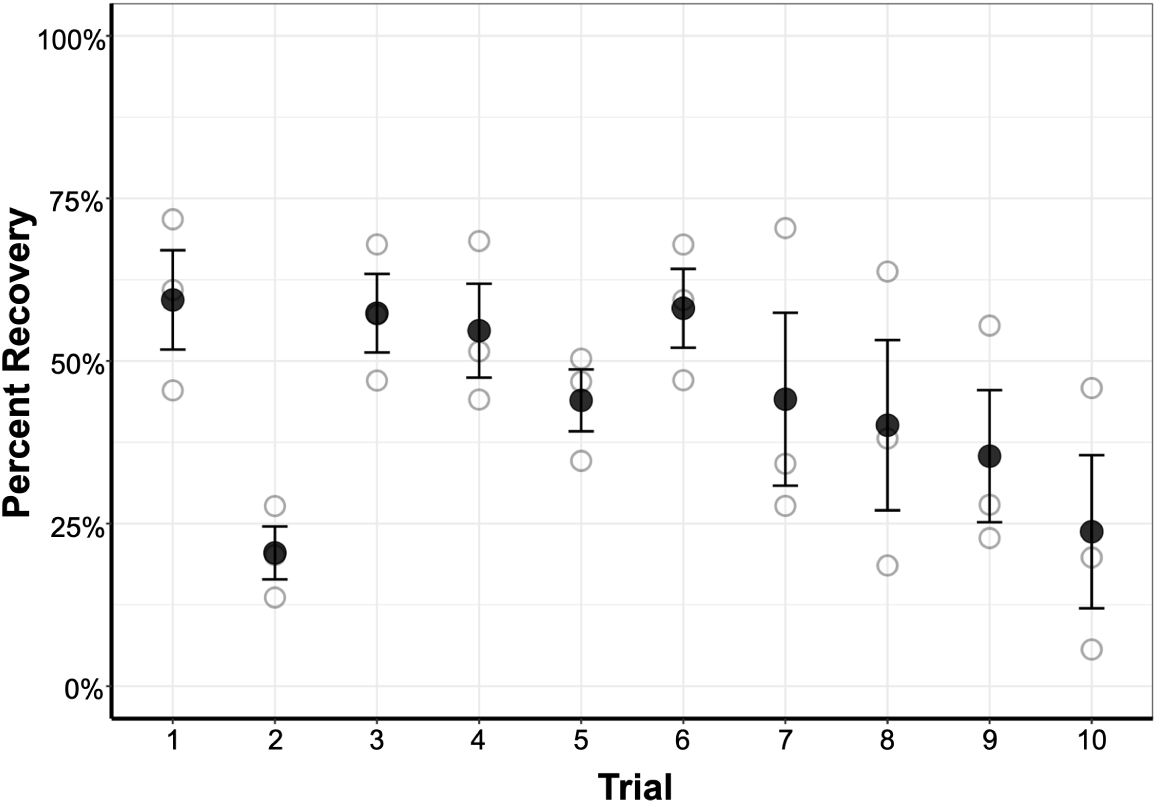
Mean recovery across the ten trials. Faded points are average values for each worm strain. Error bars show standard error of the mean.

**Figure A2.**
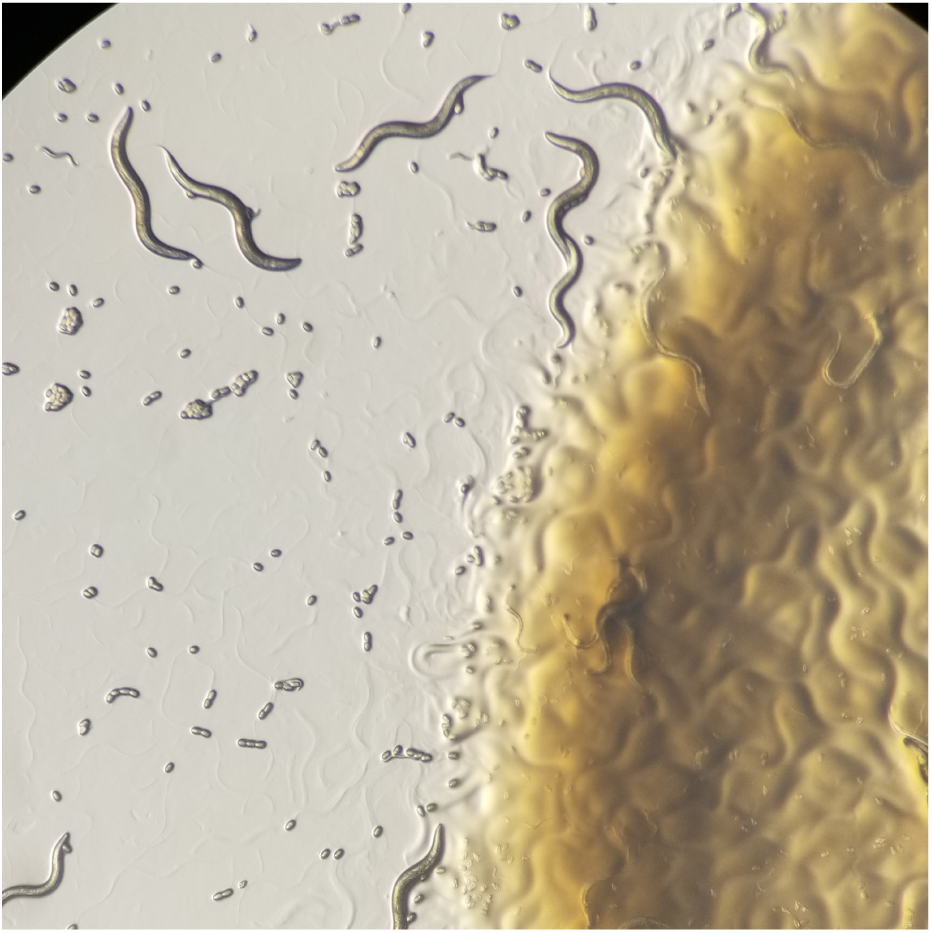
Worms of all three strains can establish populations on the beneficial bacteria.

**Table A1.** Estimated odds ratios for each value of the variables “Trial,” “Technical Replicate,” and “LB”.

| Variable | Value | Odds Ratio | 95 % CI |
| --- | --- | --- | --- |
| Trial |  |  |  |
|  | 1 | 1.00 |  |
|  | 2 | 0.18 | (0.16, 0.20) |
|  | 3 | 0.87 | (0.76, 1.00) |
|  | 4 | 0.84 | (0.74, 0.97) |
|  | 5 | 0.57 | (0.49, 0.66) |
|  | 6 | 1.03 | (0.89, 1.19) |
|  | 7 | 0.57 | (0.50, 0.66) |
|  | 8 | 0.49 | (0.43, 0.57) |
|  | 9 | 0.36 | (0.31, 0.42) |
|  | 10 | 0.20 | (0.17, 0.23) |
| Technical Replicate |  |  |  |
|  | 1 | 1.00 |  |
|  | 2 | 0.96 | (0.84, 1.10) |
|  | 3 | 1.02 | (0.89, 1.17) |
|  | 4 | 0.88 | (0.77, 1.01) |
|  | 5 | 0.91 | (0.80, 1.04) |
|  | 6 | 0.89 | (0.77, 1.02) |
|  | 7 | 0.89 | (0.77, 1.01) |
|  | 8 | 0.84 | (0.73, 0.96) |
|  | 9 | 0.88 | (0.77, 1.01) |
|  | 10 | 0.78 | (0.68, 0.90) |
| LB |  |  |  |
| | per 100 μL | 1.06 | (1.02, 1.10) |

**Table A2.**
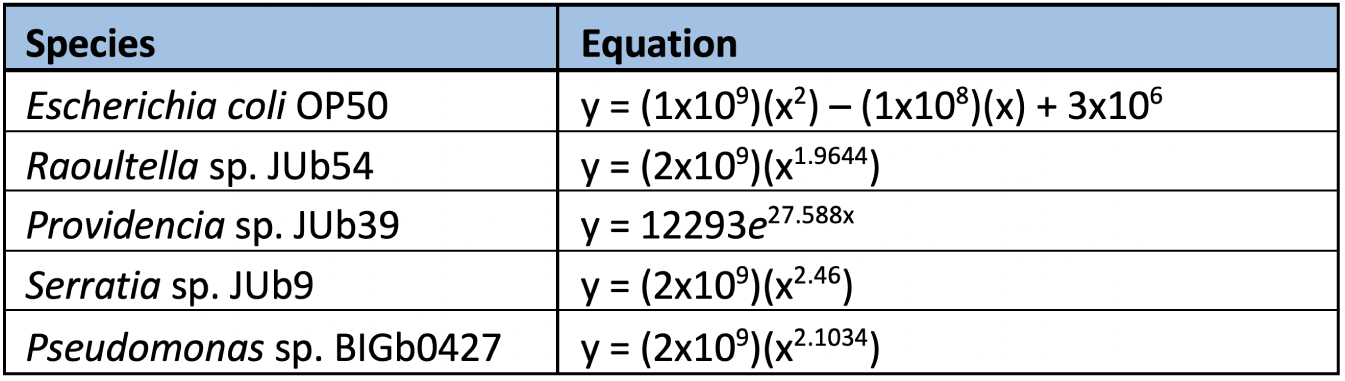
Equations used to convert absorbance to bacterial density where *x* is the absorbance and *y* is CFU/mL.

**Table A3.**
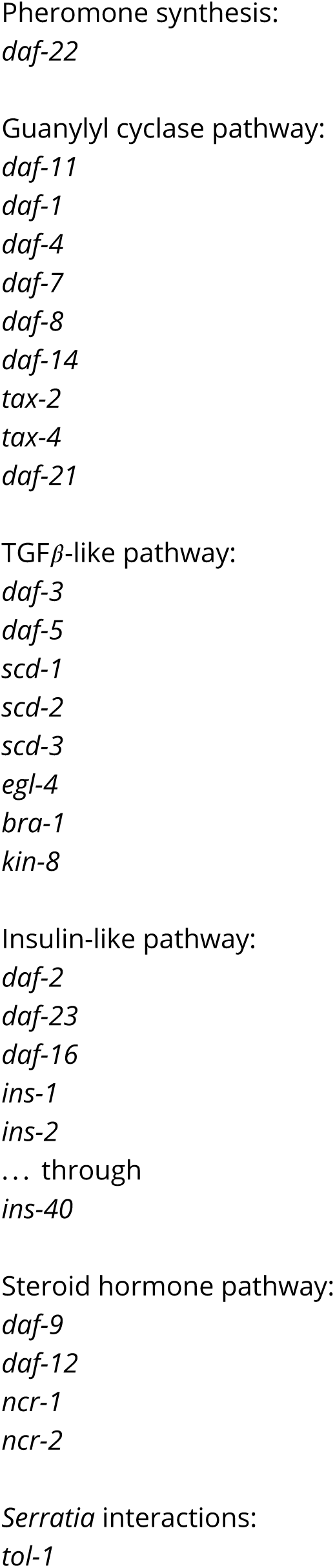
*C. elegans* dauer genes.

**Table A4.**
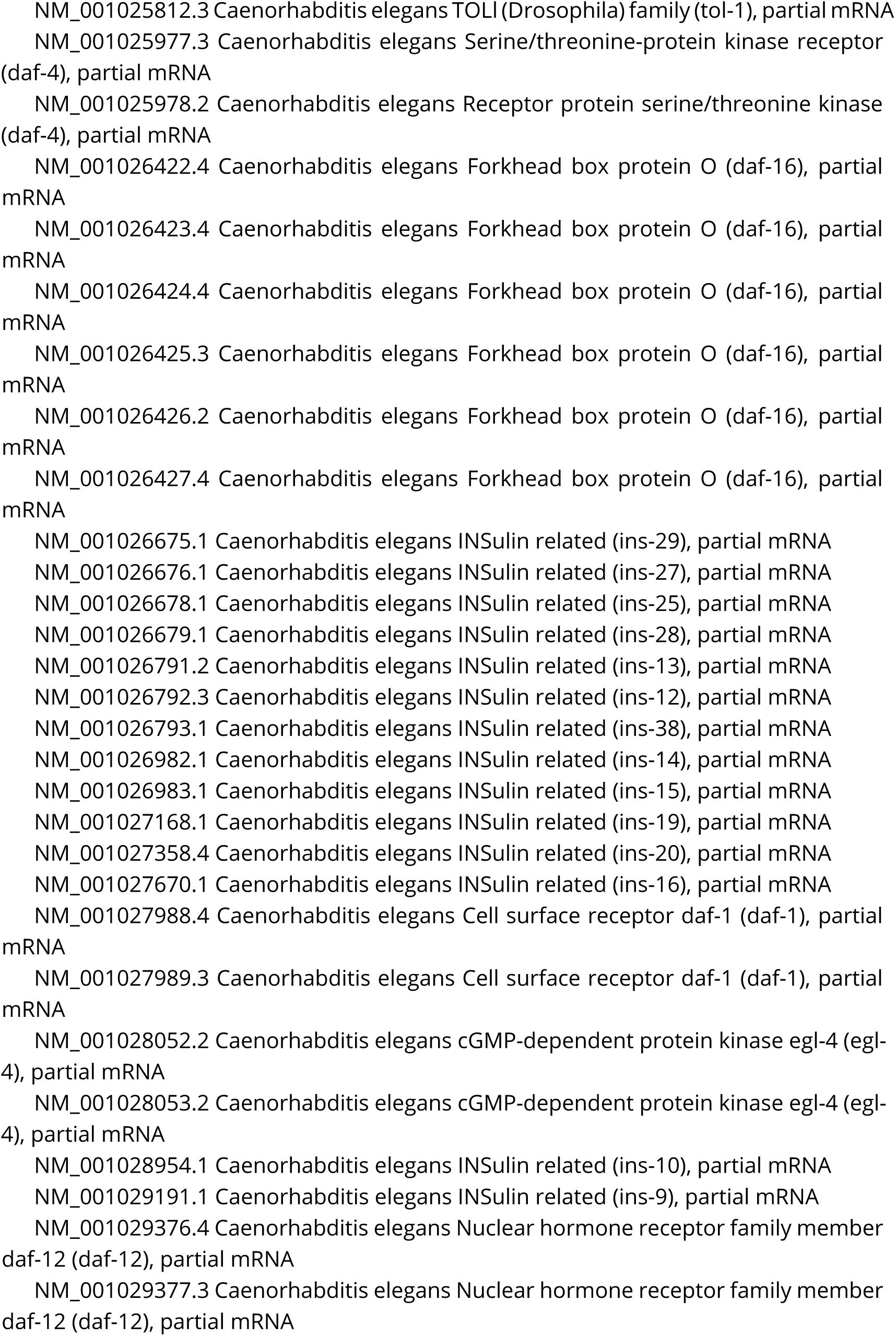
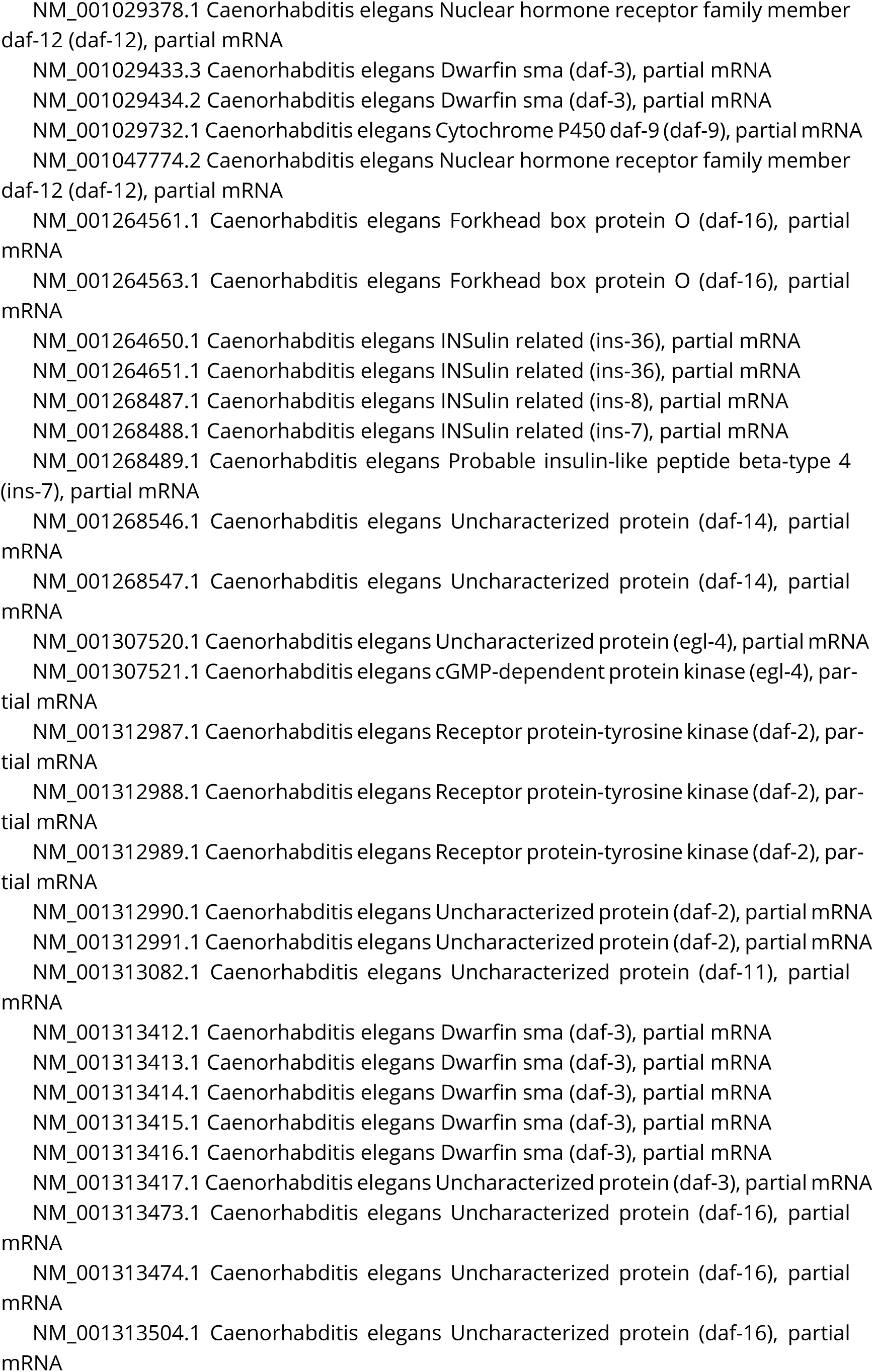
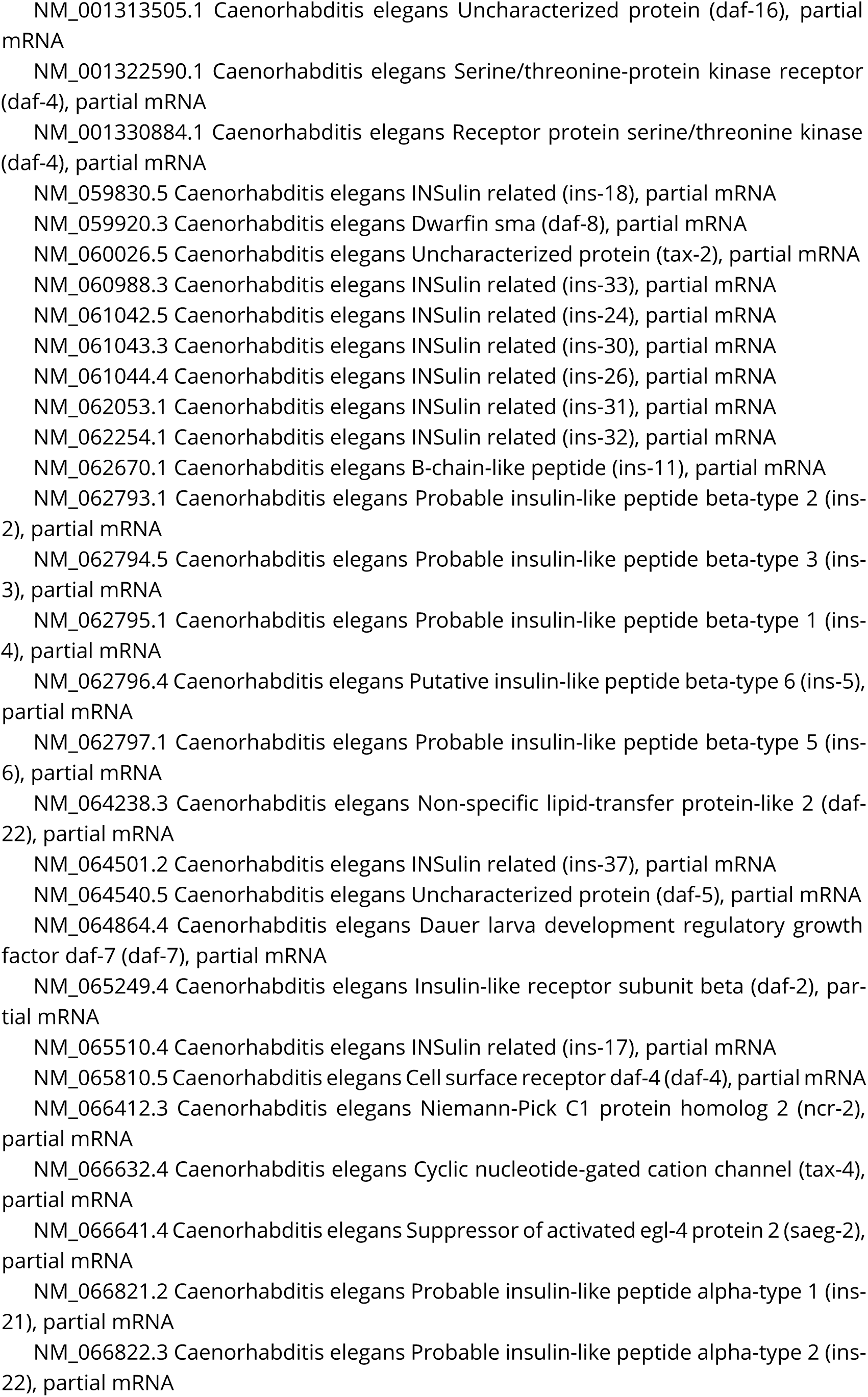
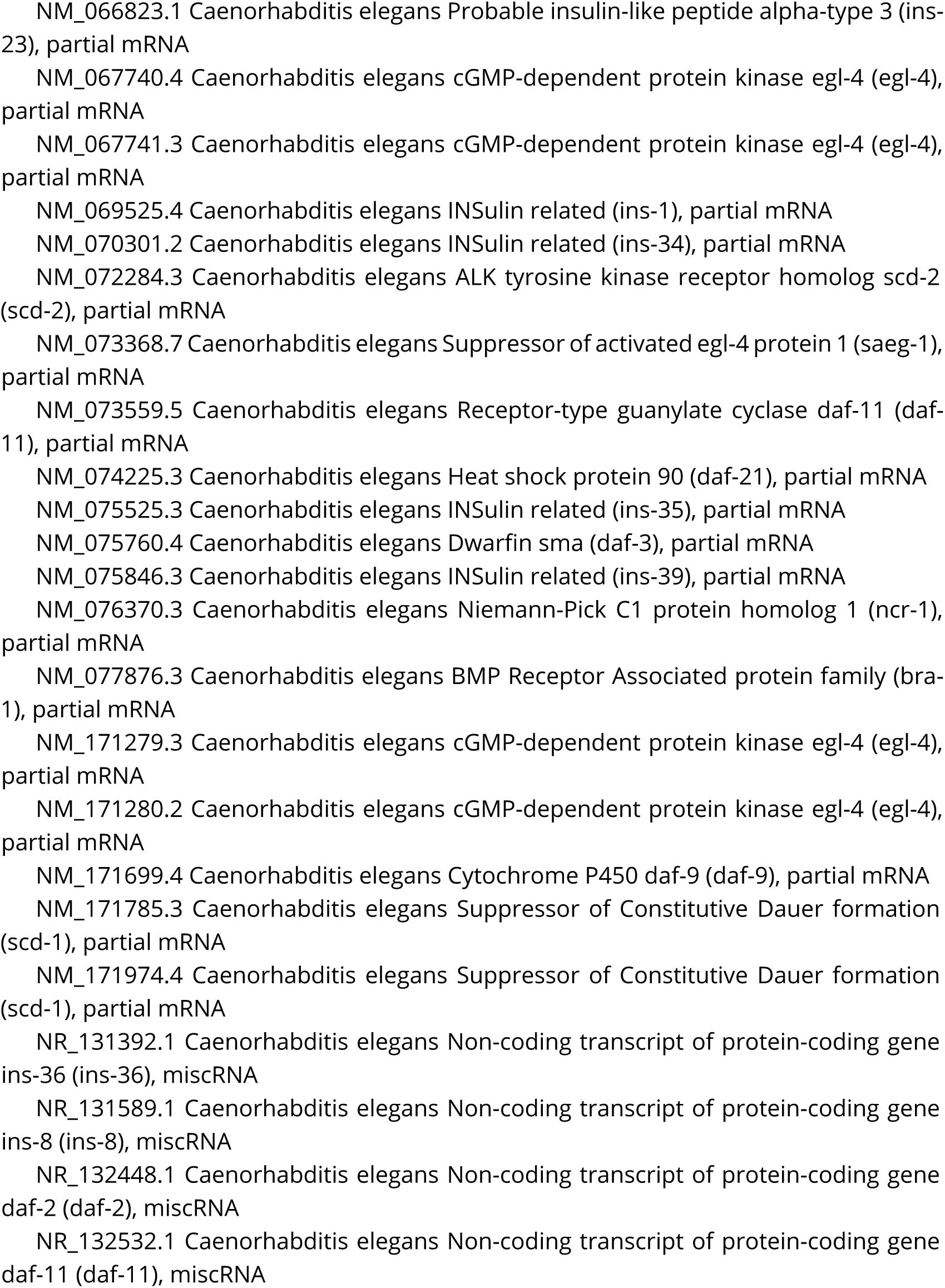
*C. elegans* dauer gene transcripts.

**Table A5.**
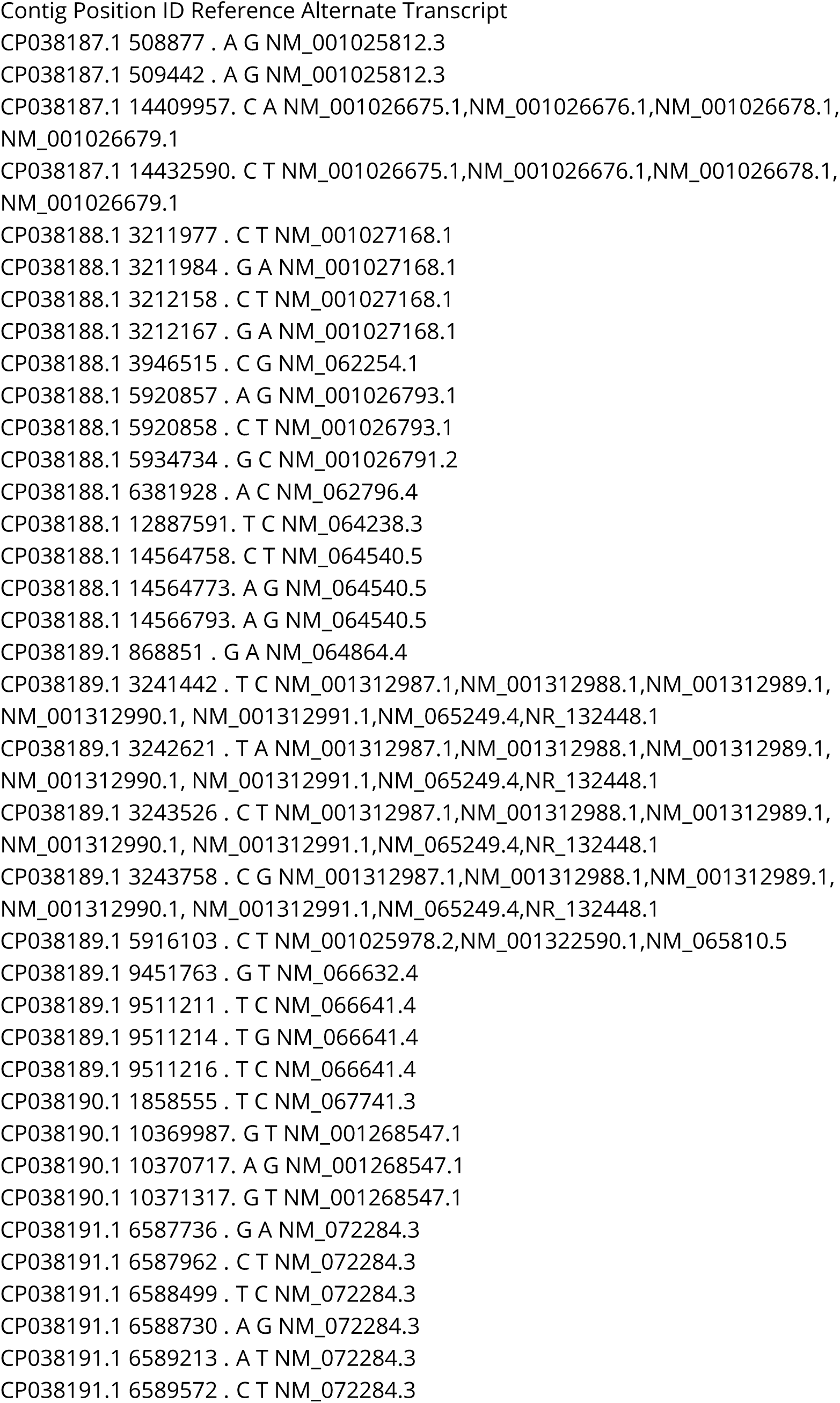
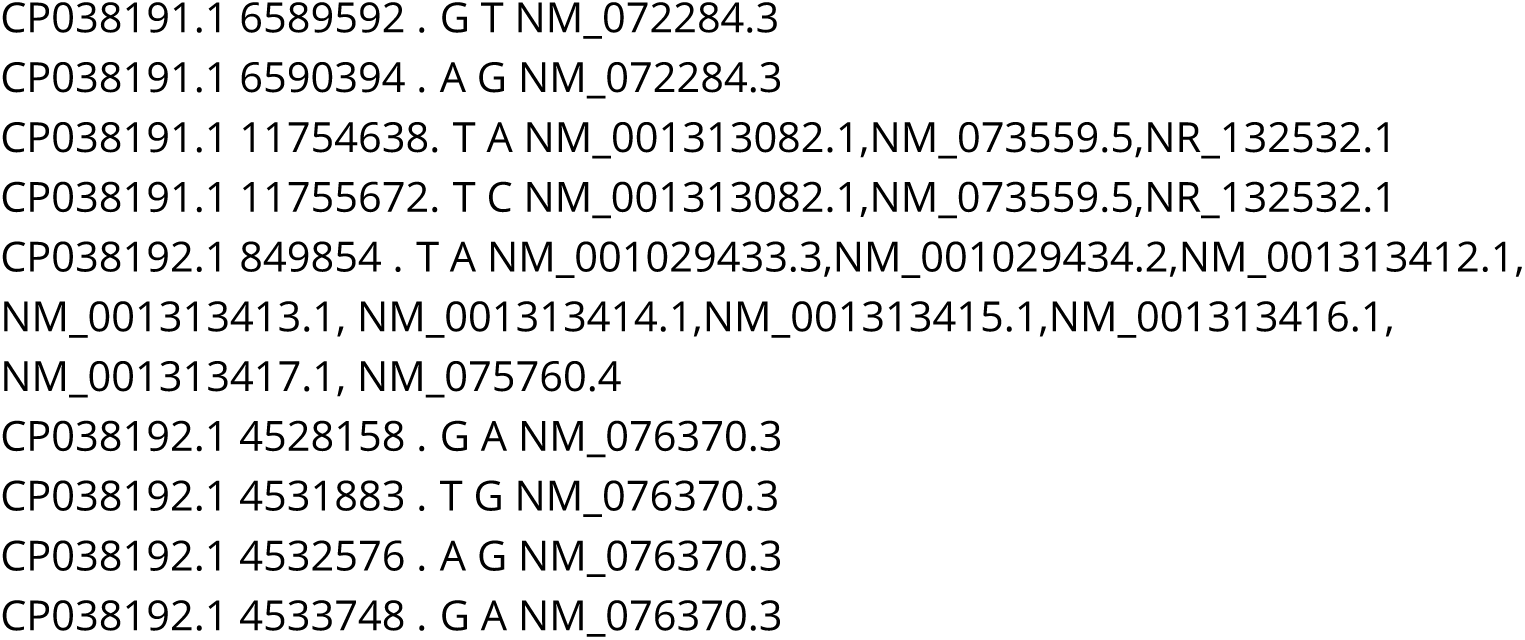
*C. elegans* CB4856 dauer transcript polymorphisms.

**Table S6.**
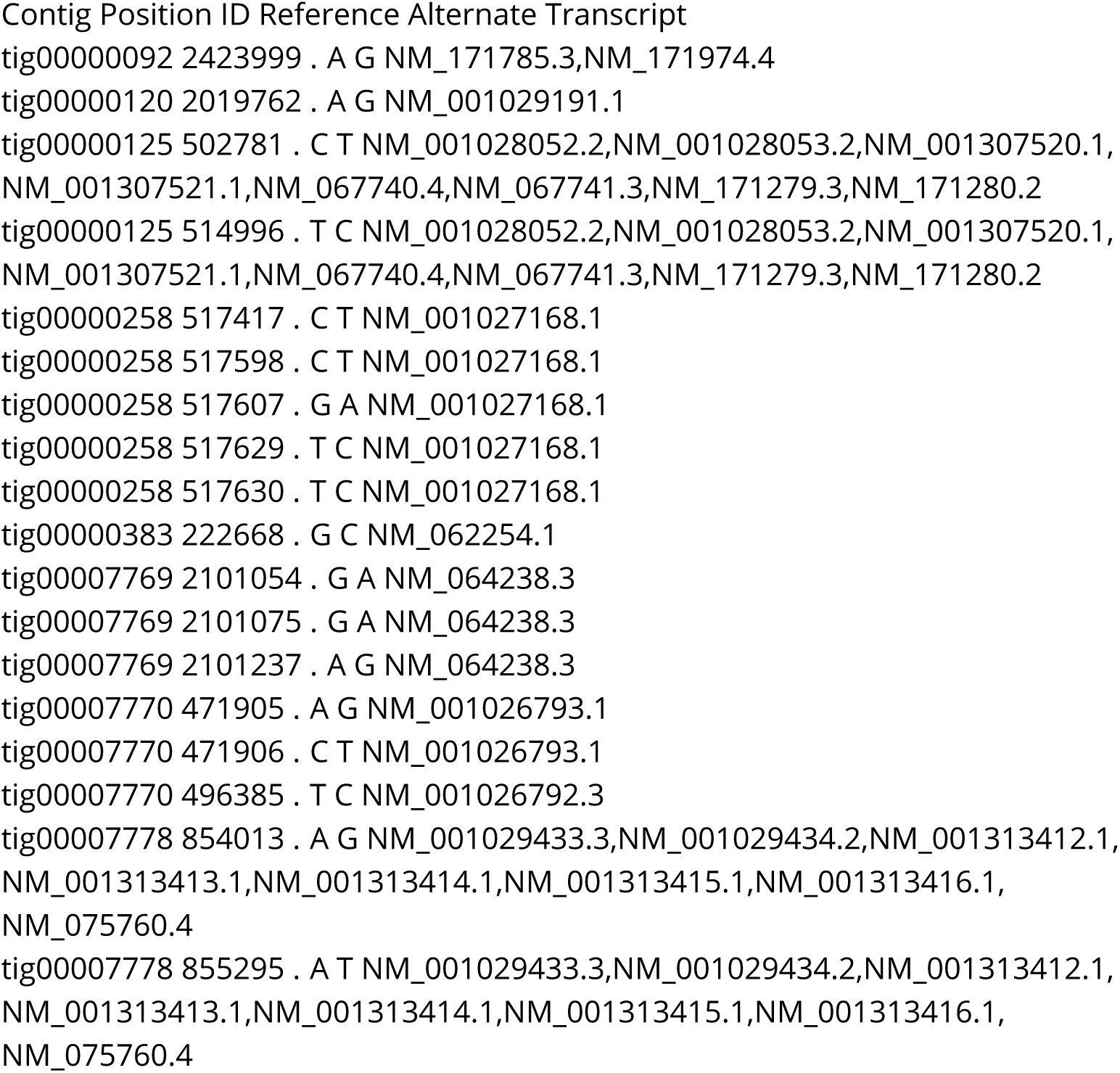
*C. elegans* JU1395 dauer transcript polymorphisms.

## Notes

#### Summary of Updates

Microbial sequencing section updated with revised names

